# A New Approach to Modeling LGMDR1: Pyrvinium-Treated *capn3b* crispant Zebrafish

**DOI:** 10.1101/2025.11.25.690358

**Authors:** Ruiz-Roldán Cristina, Valls Andrea, Immanuel Jenita, De Santis Flavia, Fernández-Torrón Roberto, López de Munain Adolfo, Saenz Amets

## Abstract

The lack of an accurate animal model for LGMDR1 is a major obstacle to therapeutic development. While murine models do not replicate the human gene expression profile, zebrafish offers a promising alternative. We generated a *capn3b* mutant zebrafish, which showed minimal phenotypic changes. However, when this model was treated with pyrvinium, a Wnt signaling inhibitor, its gene expression patterns mimic those observed in LGMDR1 patients, reinforcing the role of the Wnt pathway in LGMDR1 pathology. This pharmacologically enhanced model, despite lacking a clear phenotype in young larvae, could serve as a valuable tool for identifying potential therapeutic targets upon further investigation and validation.

## INTRODUCTION

Limb-girdle muscular dystrophy R1 calpain 3-related (LGMDR1) is an autosomal recessive muscular dystrophy due to mutations in *CAPN3* that causes progressive degeneration of the proximal muscles. For most patients, onset occurs in the second decade of life and as muscle degeneration progresses; patients require a wheelchair for mobility after around 20 years of disease progression [1, 2].

CAPN3 is a muscle-specific protease [3] whose function in muscle is not completely understood. However, it has been reported that a lack of CAPN3 alters several structures, gene expression and signaling pathways in the skeletal muscle [4–7].

To advance the development of effective treatments for any disease, a biologically relevant *in vivo* model is essential. However, finding an animal model that perfectly mimics the clinical features of calpainopathies remains elusive.

Previous attempts have used various murine models to approximate the human condition and study the effects of *CAPN3* gene alterations. However, none have fully replicated the human phenotype [4, 8–11]. Even though DNA microarray analyses were performed to establish expression profiles in mouse models, unfortunately, these failed to reflect the gene expression patterns seen in human muscle [5, 10, 12].

This difficulty is also noticeable in zebrafish, which have two *CAPN3* homologs, *capn3a* and *capn3b*, although only *capn3b* is expressed in muscle. Earlier studies on a *capn3b*-deficient zebrafish model focused on its role in other processes, such as the calpain3 involvement in the nucleolar degradation pathway of p53, liver regeneration, and autoimmune responses [13–15]. More recently, RNA-less and partial deletion *capn3b* mutants were generated using CRISPR/Cas9 mutagenesis. While this model showed no significant muscular dystrophy phenotypes under normal conditions, it was more susceptible to muscle damage under challenging conditions, such as in mechanical stress situations [16].

It is known that a correct muscle fibre function requires a delicate balance between muscle protein synthesis and proteolysis by means of the Akt/mTOR and the Wnt/β-catenin signaling pathway, among others [17–19]. Both signaling pathways are altered in LGMDR1 patient’s muscles. This could be due to the overexpression of FRZB, a Wnt signalling pathway inhibitor, and to the reduction of mTOR and downstream kinases phosphorylation in LGMDR1 patients [5, 6, 20].

We generated a new *capn3b* crispant that, as the previous model, did not accurately reflect the phenotypic characteristics of the disease. To solve this problem, we have developed a pharmacologically enhanced zebrafish model, specifically designed to mimic an impaired signaling described in patients. Therefore, we focused on the Wnt signaling pathway inhibition by means of pyrvinium. When this zebrafish model was treated with pyrvinium, the gene expression deregulation demonstrated a greater resemblance to the deregulated patterns observed in LGMDR1 patients’ muscles. This finding suggests that modulation of the Wnt signaling pathway may represent a promising therapeutic target for LGMDR1.

## MATERIALS AND METHODS

### Zebrafish Husbandry and Breeding

Adult wild-type AB strain zebrafish were maintained at 28.5 ± 1ºC in a 14 h light/10 h dark cycle in a closed recirculating tank system, according to standard protocols [21].

### Generation of Crispants

Wild-type or Casper one-cell-stage embryos were injected following published protocols [22, 23] with a mix containing the Cas9 and sgRNAs targeting the coding sequence of the gene of interest or the Cas9 and a control sgRNA (scramble) (Table S1).

### Mutation rates quantification

Genomic DNA was extracted from ten individual embryos at the stage of 24 hpf. The target region was amplified by PCR employing the primers capn3b-fw and capn3b-rev. The percentage of mutated alleles present in the PCR was quantified using the Synthego ICE analysis tool upon sequencing of the resulting amplicons.

### Treatment

Pyrvinium pamoate salt hydrate was reconstituted in Dimethyl Sulfoxide (DMSO) to a stock concentration of 1mM and diluted in E3 1X medium to a final concentration of 0.400, 0.200, 0.100, 0.050, and 0.025 μM. 0.1% DMSO in E3 1X medium. 48 hpf larvae injected with scrambled sgRNA and capn3b sgRNAs were treated with 0.1% DMSO and with each of the Pyrvinium concentrations, they were maintained at 28.5ºC with a 14:10 hour light:dark cycle. The medium was renewed at 72, 96, and 120 hpf.

### Morphological Analysis: birefringence

Larvae were anesthetized with tricaine 0.28 mg/mL, and images were captured using a stereomicroscope. After the imaging, larvae were euthanized and the birefringence was quantified as follows in Berger et al 2012 [24].

### Analysis of locomotor activity

For the locomotion analysis, the standardized 25 min dark/light cycles paradigm (5 min darkness—5 min light up to 25 min) was used, the employed methodology was based on the protocol described in Locubiche *et al*., 2024 [23].

### Statistical analysis

Statistical analysis, from locomotor activity, were performed using One way ANOVA followed by Sidak’s test.

For the study of the gene expression levels, data were expressed as the mean ± standard error of the mean (SEM) of three technical replicates. Statistical significance was determined using unpaired T-test, and a p-value < 0.05 was considered statistically significant. All computational analyses were performed using GraphPad Prism.

### RNA extraction

For RNA extraction, 16 zebrafish larvae were used per condition. The PicoPureTM RNA Isolation Kit (Applied Biosystems, Thermo Fisher Scientific, Foster City, CA, EE. UU.) was used, following the manufacturer’s instructions for RNA extraction.

### RT-qPCR

The High-Capacity cDNA Reverse Transcription Kit (Life Technologies, Carlsbad, CA, EE. UU.) was used for cDNA synthesis, following the manufacturer’s instructions.

To study gene expression levels by qRT-PCR, *SYBR Green* primers (Table S2) and *TaqMan* probes, in Gene Expression Array Cards (Applied Biosystems, Thermo Fisher Scientific, Waltham, MA, EE. UU.) (Table S3) were used. Genes were selected according to LGMDR1 patients’ expression profiling [6], however since less zebrafish probes are available, additional muscular genes were included. The CFX384 C1000 thermocycler (Bio-Rad Laboratories, Hercules, CA, EE. UU.) and QuantStudio 12K Flex Real-Time PCR System (Applied Biosystems, Thermo Fisher Scientific, Waltham, MA, EE. UU.) were used. The data were analysed using the Bio-Rad CFX Maestro programme and the ThermoFisher Connect Platform.

## RESULTS

### The CRISPR/Cas9 technology allows the efficient targeting of the *capn3b* gene

To achieve a high rate of somatic mutations in F0 zebrafish larvae, we developed a strategy using two sgRNAs targeting distinct coding regions of the *capn3b* gene. Four different sgRNA combinations were designed and tested, with their cutting efficiencies listed in Table S4. Among these, the combination of guides 2 and 3 produced the highest mutation rates across the greatest number of larvae; almost the totality of their genetic material (97.8%) contained mutations in the *capn3b* coding sequence (CDS).

### The inactivation of the *capn3b* gene and the treatment with pyrvinium do not affect locomotion

Our first step in the characterization of *capn3b* crispants was to evaluate how gene inactivation, alongside with the treatment with pyrvinium, influenced larval locomotion.

To achieve this, we tracked the swimming behavior of 144 hpf embryos. By monitoring these behavioral responses under different experimental conditions, we aimed to determine how gene knockout and drug exposure interact to affect larval motility.

We observed that all groups (scrambled and *capn3b* crispants) did not show differences in their response to the environmental illumination changes (Figure 1A).

**Figure 1.**
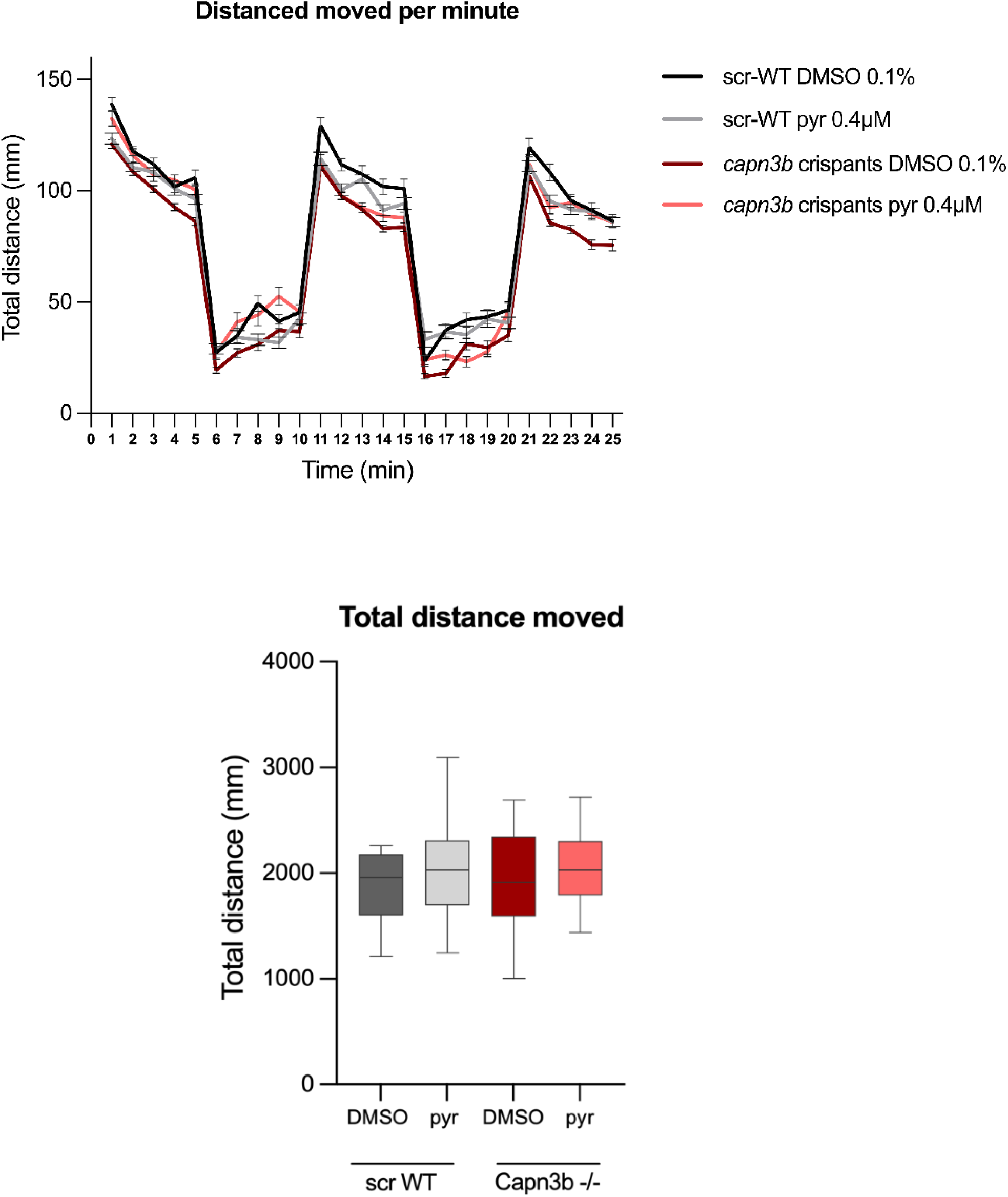
Locomotion Activity Assessment. **(A)** Total distance moved per minute during the dark/light phase. Different experimental conditions are represented in different colors as shown in the legend on the left. Error bars represent the standard error of the mean (SEM). (**B)** Total distance moved during the dark/light transition test. Scrambled is represented in gray, capn3b crispant is represented in red (0.1%DMSO) and the treated with pyrvinium 0.4 μM, in blue. Error bars represent the minimum to maximum values. All comparisons are not significant (One-way ANOVA followed by Sidak’s test for multiple comparisons).

### The inactivation of the *capn3b* gene and the treatment with pyrvinium do not affect muscular integrity

We characterized the muscular integrity in zebrafish larvae. In healthy zebrafish larvae, the sarcomere structure results in a strong birefringence signal when viewed under polarized light microscopy.

The birefringence of scrambled and *capn3b* crispants was analyzed at three developmental stages: 96, 120 and 144 hpf. We did not detect any differences between the different experimental groups, at any of the concentrations analyzed, suggesting that neither the gene KO nor the drug exposure affected the structure of muscles (Data shown only for 144 hpf, Figure S1).

### Gene expression alteration mimic the expression patterns found in LGMDR1

First of all, we found an interesting finding. We observed that *frzb* expression, highly upregulated in LGMDR1 patients’ muscles, was not deregulated in crispant models before administering pyrvinium. Therefore, in our untreated crispant model, as the Wnt signaling pathway inhibitor’s expression is not increased, this pathway is not inactivated contrary to what is observed in LGMDR1 patients (Figure 2).

**Figure 2.**
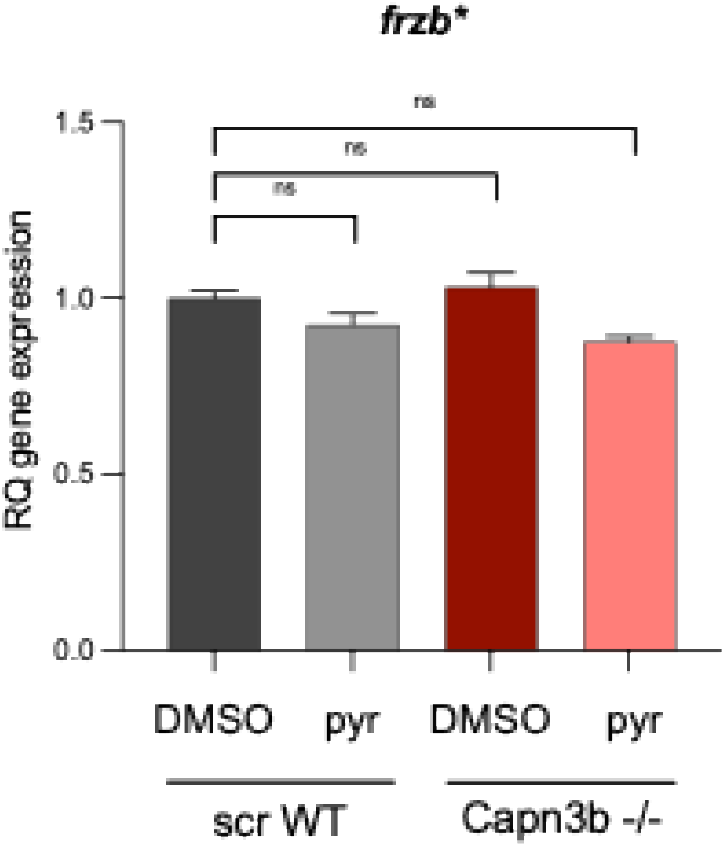
Relative expression levels of *frzb* in *capn3b*-/- model and after treatment with pyrvinium. Gene expression was analysed in both untreated control and 0.4 µM Pyrvinium-treated *capn3b*-/- zebrafish samples. Fold change threshold ≥1. Data is normalized to a reference gene, *ef1*α. *TaqMan probe. Error bars represent the minimum to maximum values. Unpaired T-Test was performed to assess statistical significance (*p-value<0.05; ***p-value<0.001 ****p-value<0.0001).

After analysing gene expression, we were able to identify that some of the genes deregulated in the muscle of LGMDR1 patients [5] were also slightly altered in the *capn3b* crispants model. Furthermore, this alteration was increased in the *capn3b* crispants model after treatment with 0.4 µM Pyrvinium, making it more similar to the alterations observed in patients.

There are genes that showed a notably similar profile in the pyrvinium treated *capn3b* crispants model, compared to LGMDR1 patients. Genes that are part of the extracellular matrix, *aspn, col1a1b, col15a1b*, were upregulated in the treated model. In contrast, genes coding for transcription factors (*fos, junbb, jun, jdp2b*) except for *trf1e*, showed significantly reduced expression. Finally, genes encoding for proteins involved in endocytosis (*vldlr*) and cell adhesion (*cd9*) were also upregulated following pyrvinium treatment (Figure 3). Vldlr is a receptor that, by binding to different ligands, participates in lipid metabolism, in a variety of biological functions and acts as a negative regulator of the Wnt signaling pathway [25–28].

**Figure 3.**
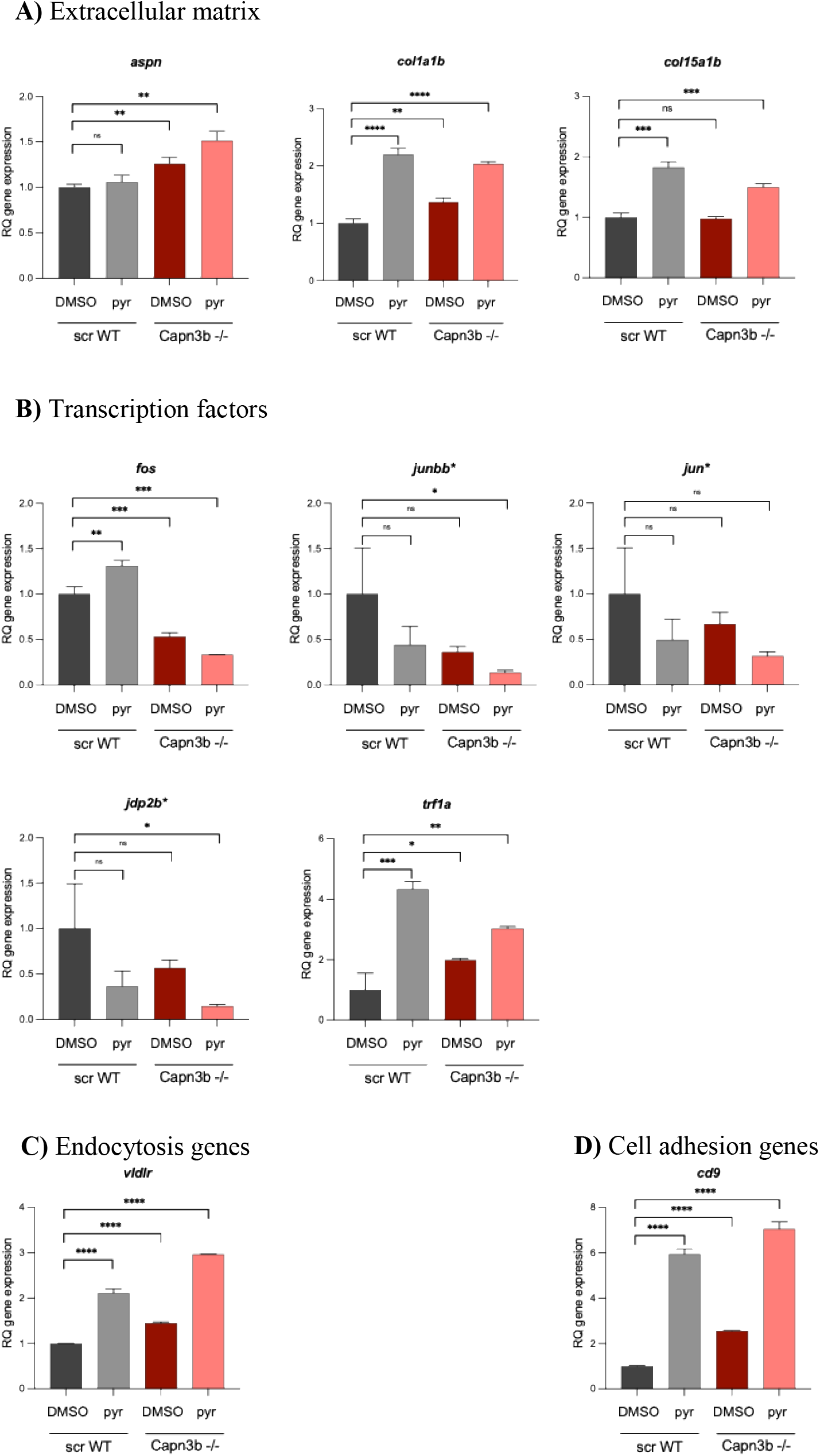
Deregulated genes in *capn3b*-/- model and after treatment with pyrvinium. Data shows the relative expression levels of **A)** Extracellular matrix (*aspn, col1a1b, col15a1b*), **B)** Transcription factors (*fos, junbb, jun, jdp2b, trf1a*), **C)** Cell adhesion genes (*vldlr*) and **D)** Endocytosis genes (*cd9*). Gene expression was analysed in both untreated control and 0.4 µM Pyrvinium-treated *capn3b*-/- zebrafish samples. Fold change threshold ≥1.5 or ≤0.5. Data is normalized to a reference gene, *ef1*α. *TaqMan probes. Error bars represent the minimum to maximum values. Unpaired T-Test was performed to assess statistical significance (*p-value<0.05; **p-value<0.01; ***p-value<0.001 ****p-value<0.0001).

The *cd9* gene encodes a cell surface glycoprotein involved in many cellular processes including differentiation, cell–cell fusion events and signal transduction [29, 30]. Additionaly, it has been described in eosinophils’ surface [31, 32].

It is worth noting that other genes, deregulated in LGMDR1 patients, showed a similar tendency in their expression, but they did not reach the established threshold of 0.5 fold-change value in the pyrvinium treated knockout model (Figure S2).

## DISCUSSION

The generated disease model, with respect to its muscular morphology and function, does not resemble the characteristics observed in LGMDR1 patients. However, it is noteworthy that this model is of great interest due to the rapid generation, made possible by CRISPR/Cas9 technology. This technology has been further demonstrated by other studies to generate zebrafish models for different muscular dystrophies, providing valuable insights into disease mechanisms and potential therapies [33–35].

On the one hand, the lack of *frzb* expression deregulation in the crispant model prior to the treatment, unlike what is observed in LGMDR1 patients’ muscles, suggests that Wnt signaling pathway is not inhibited in this model.

This signaling pathway is one of the most relevant for protein synthesis. Given the high protein turnover in the muscle fiber, the impairment of its activity, combined with calpain 3 deficiency, poses an additional limitation for the proper muscle function in LGMDR1 patients. Nevertheless, in our zebrafish model, since the Wnt signaling does not appear to be inhibited, it could be suggested that its maintained activity might enable the model to avoid the dystrophic phenotype. However, further studies are required to confirm this hypothesis.

Based on the alterations in gene expression following pyrvinium treatment, relevant similarities to the LGMDR1 pathophysiology were observed. Fibrosis, the excessive accumulation of connective tissue that replaces the muscle fibers is a hallmark of muscular dystrophies, leading to impaired function and reduces muscular strength. Notably, the increase in collagen expression is not observed in the untreated crispant model. However, the Wnt signaling inhibition induced by drug administration elevates the expression of two types of collagens (*col1a1b, col15a1b*), which are also upregulated in LGMDR1 patients [5]. This model could provide a fibrosis-presenting system for future studies.

Regarding the expression of the transcription factors genes, it is important to note that, although their expression is already reduced in the untreated crispant model, this reduction is even more pronounced following drug administration. This finding is particularly relevant, as the strongest downregulation of the transcription factors expression in the patients’ muscles is observed along with the most homogeneous downregulation pattern across the biological groups [5].

Finally, among the deregulated genes with distinc functions, it has been reported that the very-low-density lipoprotein receptor (VLDLR) heterodimerizes with LRP6 and that this interaction may represent a novel mechanism for regulating Wnt/β-catenin signaling. Notably, VLDLR negatively regulates Wnt signaling pathway [28, 36, 37]. This observation aligns with the finding in patients and represents an interesting mechanism that require further investigation.

The fact that cd9 is present on the eosinophils surface [31, 32] and that its expression is significantly increased in the treated model is of great interest, especially since eosinophilic infiltration has been identified in LGMDR1 younger patients [38, 39]. This may suggest that, since these zebrafish models are still too young, they could replicate observations seen in younger patients. While it is true that we cannot definitively demonstrate the presence of eosinophilia, it has been reported a conserved role for eosinophils in the zebrafish [40].

Therefore, the results obtained in the zebrafish model treated with pyrvinium, an inhibitor of the Wnt signalling pathway, support the relevance of Wnt pathway regulation in disease progression.

On the one hand, the absence of clinical or functional characteristics similar to those of LGMDR1 patients makes this model not fully validated for drug evaluation. On the other hand, the improved similarity in gene expression patterns achieved with the drug Pyrvinium paves the way to obtain a potential animal model [18, 41].

However, it is important to note that due to animal experimentation limitations, the study only reached the larval stage at 144 hpf. At this early stage, the fish may not yet exhibit a clear phenotype due to their age [42]. Ideally, to better mimic the clinical onset of the disease in patients [1, 2] a more advanced stage should be reached, such as the juvenile stage (approximately 30 days old) or later. Evaluating these fish at a more mature stage would help to determine if we have generated a truly representative disease model. Therefore, we suggest that subsequent studies are required on juvenile and adult models to confirm the possible phenotype.

## Author Contributions

Conceptualization, A.S.; Methodology, C.R.-R., and A.S.; Validation, C.R.-R, J.I., and A.S.; Formal analysis, A.V., C.R.-R., and A.S.; Investigation, A.V., C.R.-R., J.I. and A.S.; Resources, A.S.; Writing—original draft, C.R.-R and A.S.; Writing—review & editing, C.R.-R., A.V., J.I., F.DS, R.F-T, A.L.d.M. and A.S.; Visualization, A.S.; Supervision, A.S.; Project administration, A.S.; Funding acquisition, A.S. All authors have read and agreed to the published version of the manuscript.

## Institutional Review Board Statement

For breeding of zebrafish, ethical approval was obtained from Generalitat de Catalunya “Direcció General de Polítiques Ambientals i Medi Natural” under referential procedure number 18-011-ISA-RESOLUCIO 9421: Breeding of zebrafish (*Danio rerio*) for the maintenance of lines and the creation of transgenics intended to study human diseases.

## Acknowledgments

We acknowledge Ana García Durán for her valuable advice and support in the design of the experimental process.

## FIGURE LEGENDS

**Figure S1.**
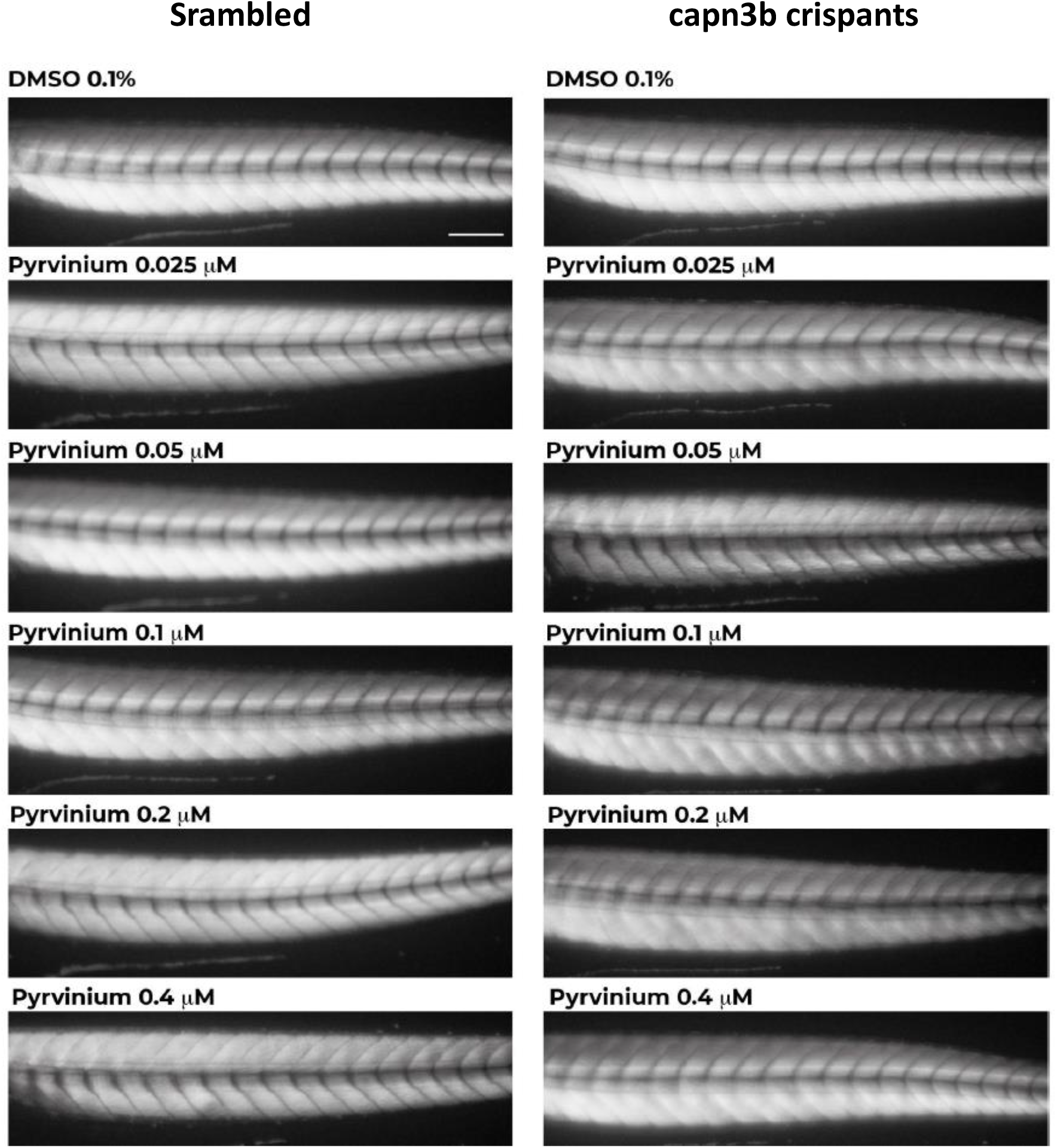
Representative pictures of the birefringence signal of scrambled (left panel) and capn3b crispants (right panel) at 144 hpf. Different concentrations of the pyrvinium treatment are shown in increasing order, from top to bottom (0.1%DMSO; pyrvinium 0.025 μM, pyrvinium 0.05 μM, pyrvinium 0.1 μM, pyrvinium 0.02 μM, pyrvinium 0.4 μM). Scale bar= 100 μm.

**Figure S2.**
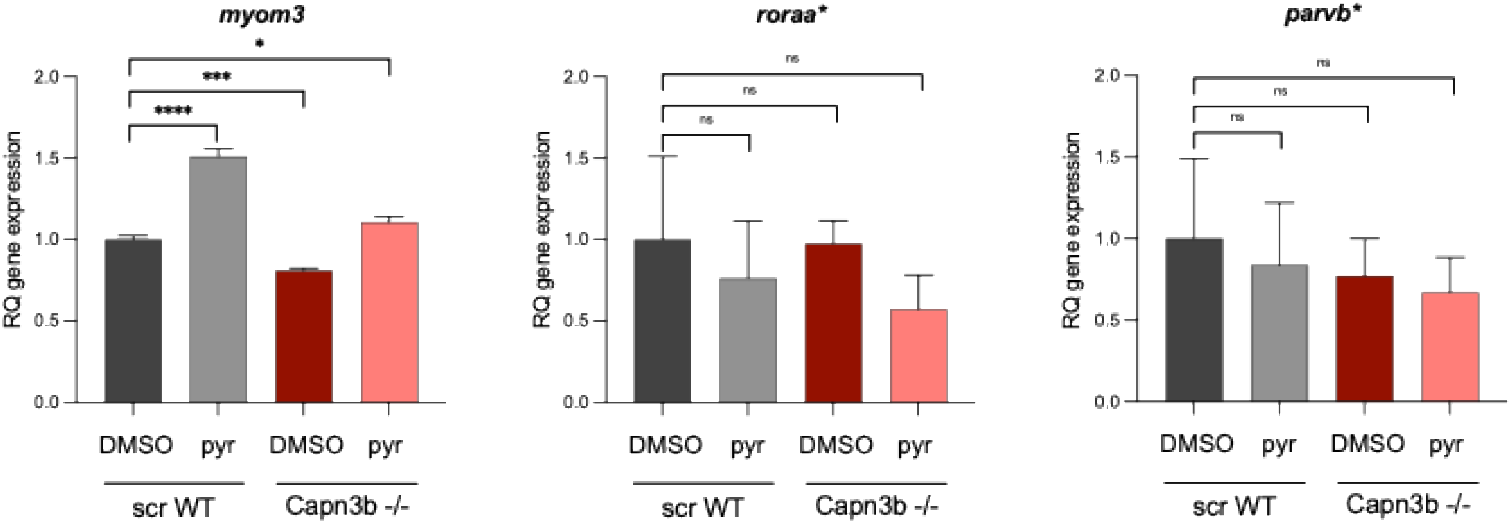
Genes with subtle alterations in expression levels in *capn3b*-/- model and after treatment with pyrvinium. Data shows the relative expression levels of *myom3, roraa* and *parvb*. Gene expression was analysed in both untreated control and 0.4 µM Pyrvinium-treated *capn3b*-/- zebrafish samples. Fold change threshold ≥1. Data is normalized to a reference gene, *ef1*α. *TaqMan probes. Error bars represent the minimum to maximum values. Unpaired T-Test was performed to assess statistical significance (*p-value<0.05; ***p-value<0.001 ****p-value<0.0001).

## SUPPLEMENTAL MATERIAL

**Table S1.**
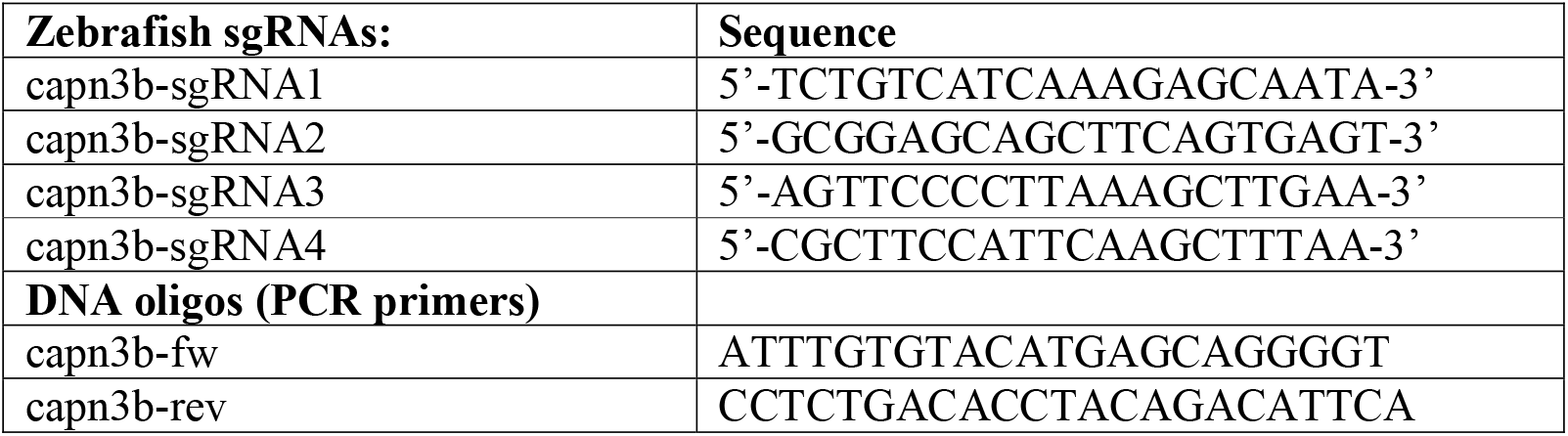
Sequences of sgRNA and the corresponding primers for *capn3b* analysis and sequencing.

**Table S2.**
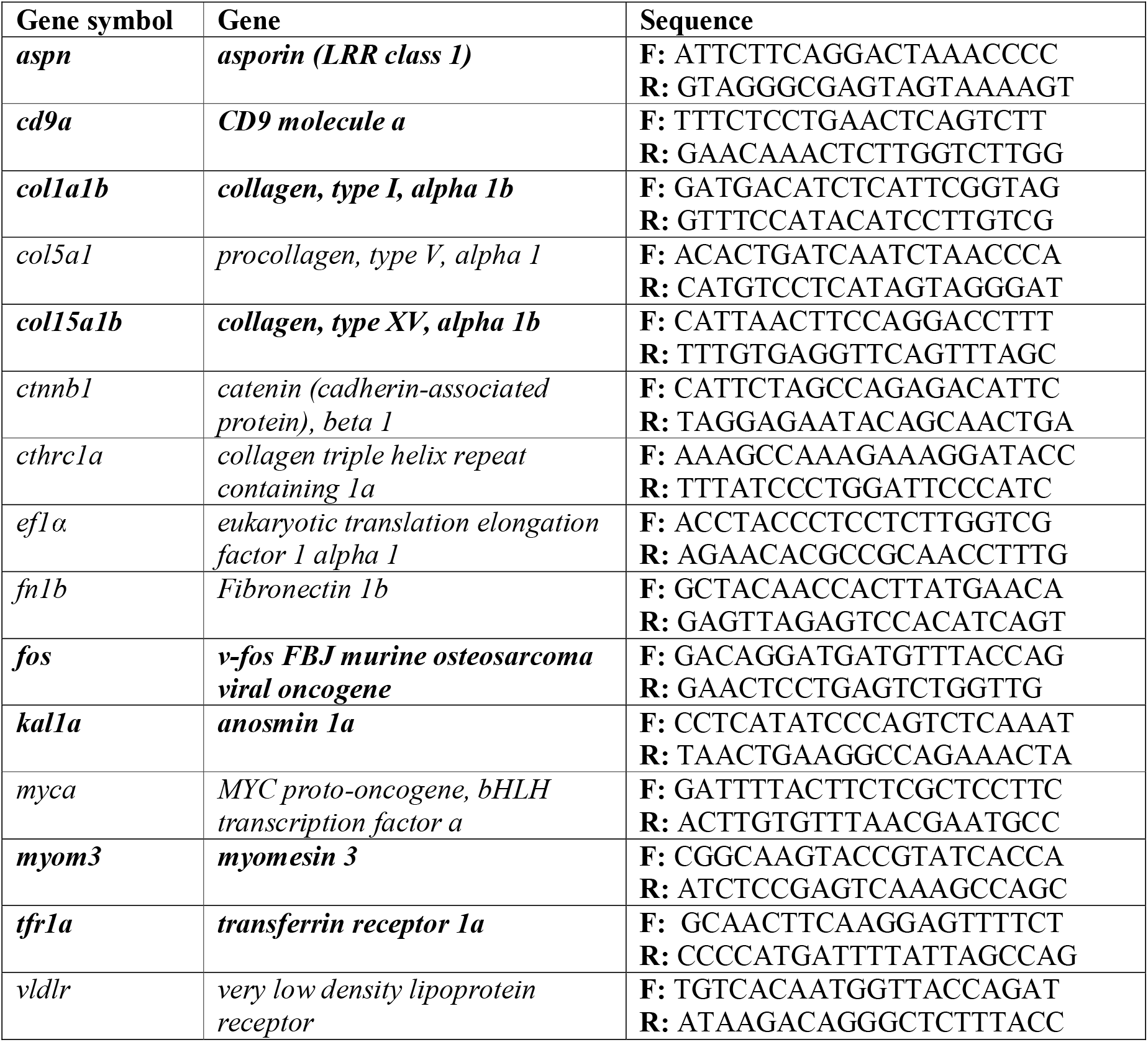
Sequences of *SYBR-Green* primers.

**Table S3.**
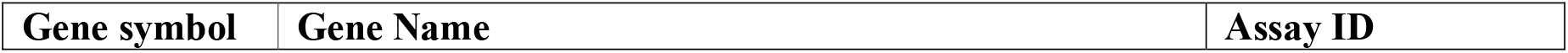
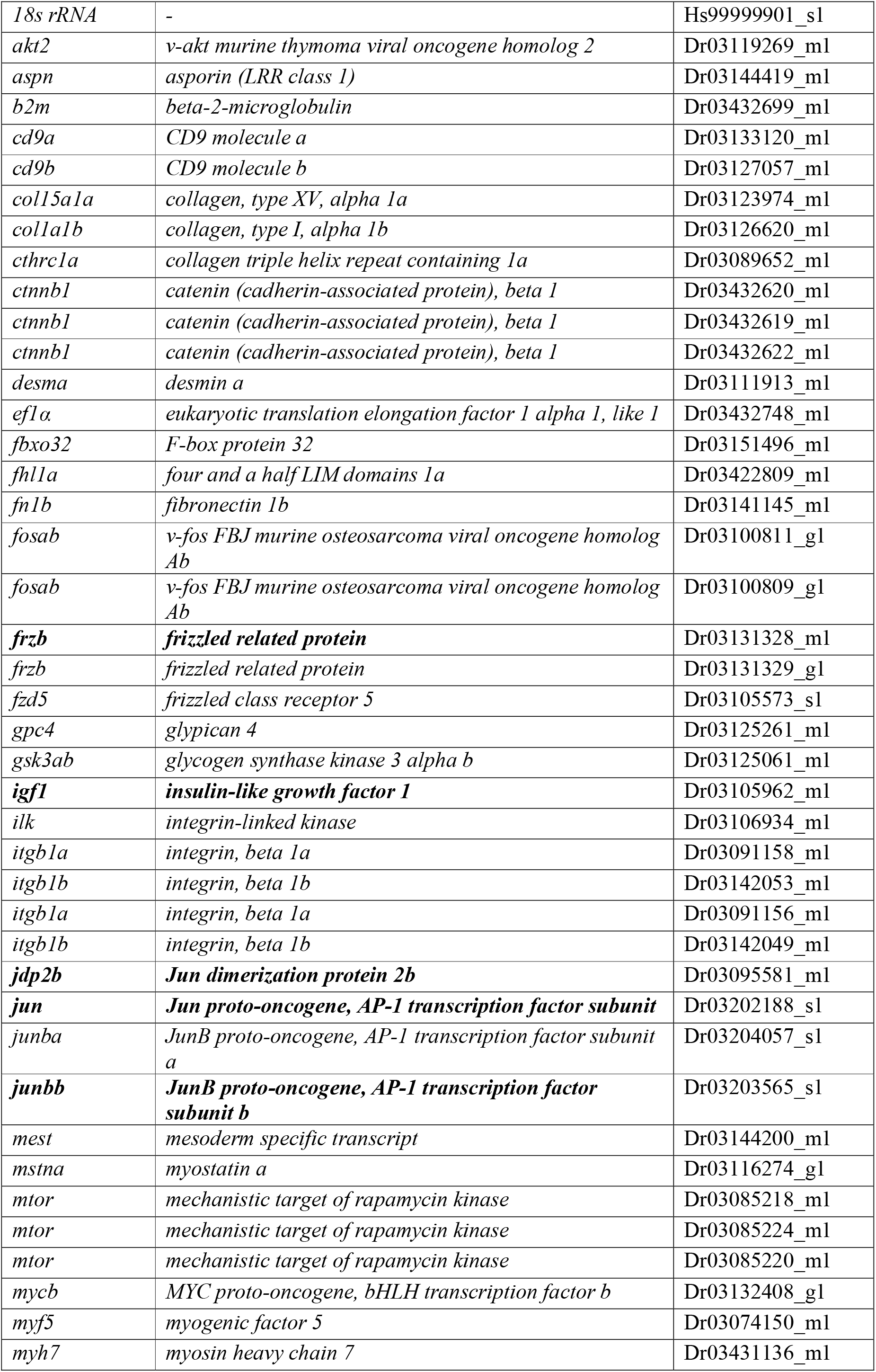
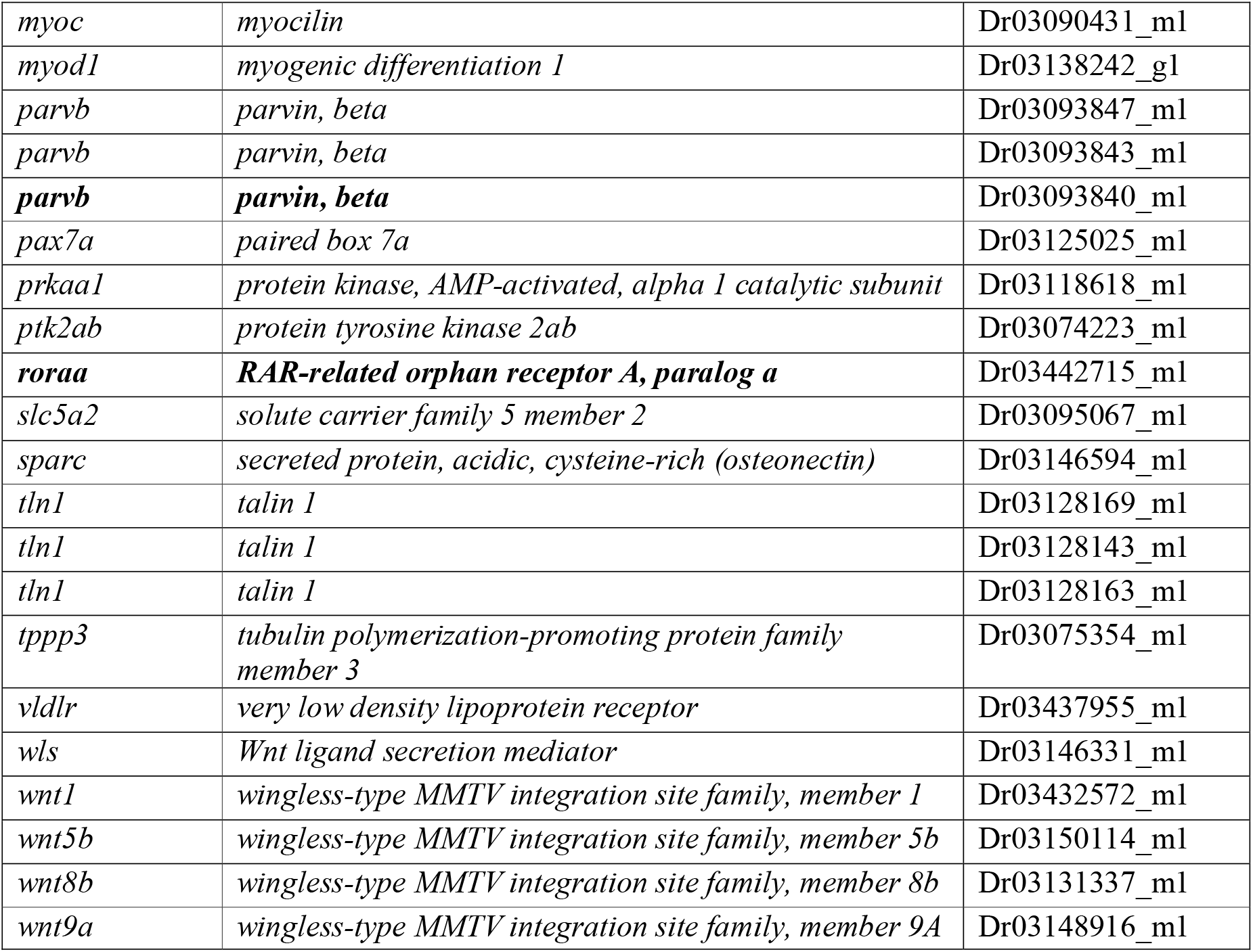
*TaqMan* probes.

**Table S4.** sgRNAs cutting efficiency (mean +/- SEM).

| *capn3b* gene | sgRNAs combinations | Efficiency |
| --- | --- | --- |
|  | 1+3 | 94 +/- 4.4 |
|  | 1+4 | 49.7 +/- 11.3 |
|  | 2+3 | 97.8 +/- 1.9 |
|  | 2+4 | 69 +/- 7.9 |

**Table S5.**
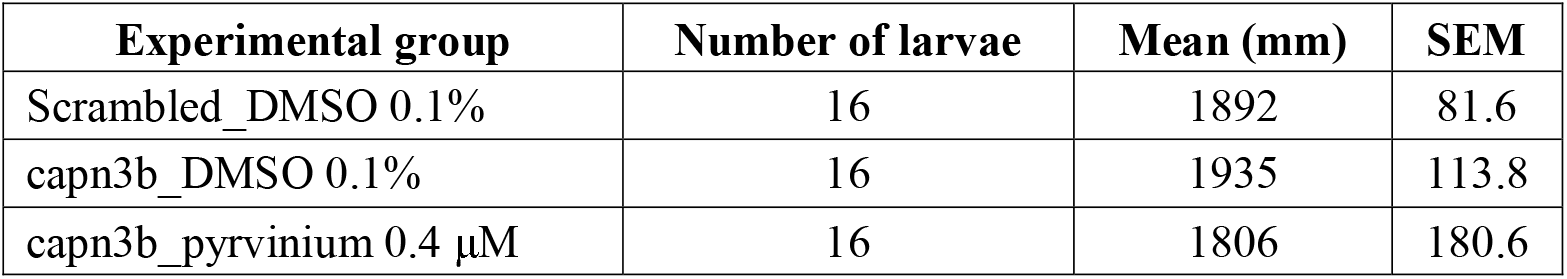
Total distance moved during the trial by each larva.

| Experimental group | Number of larvae | Mean (mm) | SEM |
| --- | --- | --- | --- |
| Scrambled_DMSO 0.1% | 16 | 1892 | 81.6 |
| capn3b_DMSO 0.1% | 16 | 1935 | 113.8 |
| capn3b_pyrvinium 0.4 μM | 16 | 1806 | 180.6 |

